# Synergistic action of the *Arabidopsis* spliceosome components PRP39a and SmD1 in promoting post-transcriptional transgene silencing

**DOI:** 10.1101/2022.08.10.503523

**Authors:** Jérémie Bazin, Emilie Elvira-Matelot, Thomas Blein, Vincent Jauvion, Nathalie Bouteiller, Jun Cao, Martin D. Crespi, Hervé Vaucheret

## Abstract

Besides regulating splicing, the conserved spliceosome component SmD1 was shown to promote posttranscriptional silencing of sense transgenes (S-PTGS). Here, we show that the conserved spliceosome component PRP39a also plays a role in S-PTGS. However, PRP39a and SmD1 actions appear distinct in both splicing and S-PTGS. Indeed, RNA-seq analysis of *prp39a* and *smd1* mutants identified different sets of deregulated mRNAs and non-coding RNAs, both at expression level and alternative splicing genome-wide. Moreover, double mutant analyses involving *prp39a* or *smd1* and RNA quality control (RQC) mutants revealed genetic interactions of SmD1 and PRP39a with distinct nuclear RQC machineries, suggesting synergistic rather than redundant roles in the RQC/PTGS interplay. Supporting this hypothesis, a *prp39a smd1* double mutant exhibited enhanced suppression of S-PTGS compared with single mutants. Because no major changes in the expression of PTGS or RQC components or in small RNA production were identified in *prp39a* and *smd1* mutants, and because *prp39a* and *smd1* mutations do not alter PTGS triggered by inverted-repeat transgenes directly producing dsRNA (IR-PTGS), PRP39a and SmD1 seem to synergistically promote a step specific to S-PTGS. We propose that, independent of their specific roles in splicing, PRP39a and SmD1 limit 3’-to-5’ and 5’-to-3’ degradation of transgene aberrant RNAs, respectively, thus favoring the export of aberrant RNAs to the cytoplasm where their transformation into dsRNA initiates S-PTGS.

## Introduction

RNA interference (RNAi) is a conserved mechanism that regulates endogenous gene expression and acts as a defense mechanism against invasive sequences of endogenous (transposons) or exogenous (virus, transgenes) origins. RNAi is initiated by double-stranded RNA (dsRNA) formed by the transcription of an inverted repeat, the copying of single-stranded RNA (ssRNA) by RNA-dependent RNA polymerases (RDRs), or convergent transcription on both strands of the DNA. These dsRNA are processed by DICER or DICER-like (DCL) proteins into microRNAs (miRNAs) or small interfering RNAs (siRNAs). These small RNAs are loaded onto ARGONAUTE (AGO) proteins to mediate sequence-specific gene silencing. Generally, miRNAs silence complementary mRNAs at the post-transcriptional level, while siRNAs can act at either transcriptional or post-transcriptional level to silence homologous endogenous genes, transgenes, transposons and viruses (Bartel, 2004; Baulcombe, 2004; Lippman and Martienssen, 2004).

In plants, post-transcriptional RNAi triggered by loci exhibiting an inverted-repeat (IR) structure is referred to as IR-PTGS. This process does not require RDR6 because dsRNAs are formed by the folding of self-complementary RNAs. In contrast, plant post-transcriptional RNAi triggered by loci that are not supposed to produce dsRNA is referred to as sense PTGS (S-PTGS). It is assumed that such loci produce a type of aberrant RNAs (abRNA) that are prone to be converted to dsRNA by RDR6. However, because RDR6 resides in cytoplasmic foci referred to as siRNA-bodies (Jouannet et al., 2012), abRNAs could only be transformed into dsRNA by RDR6 if they escape degradation by specialized RNA quality control (RQC) pathways that normally intercept and remove abRNAs in the nuclear and cytoplasmic compartments. Therefore, it is assumed that loci undergoing S-PTGS produce large amounts of abRNAs that somehow saturate the RQC machinery, thus allowing a sufficient amount of abRNAs to reach siRNA-bodies to initiate S-PTGS. Supporting this hypothesis, *Arabidopsis dpc1, dcp2, hen2, ski2, ski3, upf1, upf3, vcs, xrn3* and *xrn4* mutations, which compromise RQC at different steps, promote or enhance RDR6-dependent PTGS of both transgenes (Gazzani et al., 2004; Gy et al., 2007; Gregory et al., 2008; Thran et al., 2012; Moreno et al., 2013; Martinez de Alba et al., 2015; Yu et al., 2015) and endogenous genes (Martinez de Alba et al., 2015; Zhang et al., 2015).

Following the transformation of abRNAs into dsRNA by RDR6, DCL4 and DCL2 process dsRNAs into 21- and 22-nt siRNAs, respectively. These siRNAs form a complex with AGO1 and guide silencing of complementary mRNAs. Once S-PTGS is initiated, the biogenesis of 21- and 22-nt siRNAs usually extends along the entire length of the mRNA in a process known as transitivity (Vaistij et al., 2002; Mlotshwa et al., 2008; Parent et al., 2015). Furthermore, once induced in cells where RQC is impaired or saturated, S-PTGS moves systemically throughout the plant. The existence of systemic PTGS signals was first demonstrated by grafting non-silenced transgenic scions onto silenced transgenic rootstocks carrying the same transgene (Palauqui et al., 1997), and subsequent grafting experiments with mutant rootstocks and scions has identified several plant genes that are required for transmission of systemic PTGS from rootstocks and/or reception of systemic PTGS in scions (Brosnan et al., 2007; Taochy et al., 2017; Taochy et al., 2019). Nevertheless, full systemic S-PTGS has only been observed in the cases of transgenes, suggesting that transgenes differ from endogenous genes in a way that makes them particularly prone to trigger S-PTGS all over the plant. Therefore, it is likely that if RQC is impaired or saturated in one cell, the entry of abRNAs into siRNA-bodies and the triggering of S-PTGS remains confined to this cell and its surrounding cells in the case of endogenous genes, thus preventing the deleterious effects that could result from systemic S-PTGS of essential genes. Supporting this hypothesis, impairing RQC constitutively causes the death of plants, which can be suppressed by impairing RDR6 (Martinez de Alba et al., 2015; Zhang et al., 2015). In contrast, transgenes, which obviously are dispensable because they are not present naturally, generally are silenced systemically, suggesting that transgenes are treated like exogenous invaders such as viruses, which need to be systemically silenced all over the plant.

Genetic screens for S-PTGS-deficient mutants have been instrumental in identifying S-PTGS components, whereas genetic screens for S-PTGS-enhanced mutants have been instrumental in identifying RQC components that counteract S-PTGS. However, little is known about factors that counteract the action of RQC and favor S-PTGS. We previously reported that the conserved spliceosome component SmD1, which localizes in the nucleoplasm, promotes transgene S-PTGS, even when triggered by intron-less transgenes, and has no effect on transgene IR-PTGS (Elvira-Matelot et al., 2016). Because S-PTGS is restored in double mutants impaired in SmD1 and the RQC factor XRN3, which also localizes in the nucleoplasm, we proposed that SmD1 protects transgene abRNAs from degradation by XRN3. Thus, beside its role in splicing, a spliceosome component plays a role in the interplay between RQC and S-PTGS in the nucleus (Elvira-Matelot et al., 2016). Here, we show that another conserved spliceosome component, PRP39a, promotes S-PTGS, even when triggered by intron-less transgenes, while it has no effect on IR-PTGS. S-PTGS is restored in *prp39a hen2* and *smd1 xrn3* double mutants, but not in the *prp39a xrn3* and *smd1 hen2* double mutants, suggesting distinct interactions of PRP39a and SmD1 with the HEN2- and XRN3-dependent RQC pathways, respectively. Impairing simultaneously PRP39a and SmD1 further suppresses S-PTGS demonstrating a synergistic action of these proteins. These proteins also affect the alternative splicing of different pre-mRNA targets genome-wide. Hence, PRP39a and SMD1 define two nuclear pathways with dual roles in splicing and S-PTGS regulation.

## Results

### SGS15 encodes an ortholog of the yeast splicing factor PRP39

The Arabidopsis *2a3* line carries a transgene consisting of the *NIA2* gene under the control of the 35S promoter. This line undergoes S-PTGS of the *35S:NIA2* transgene and of the endogenous *NIA1* and *NIA2* genes in each generation, resulting in a chlorotic phenotype (Elmayan et al., 1998b). Fast-neutron mutagenesis followed by a screen for plants that remain green yield several S-PTGS-deficient mutants that have been described previously (Adenot et al., 2006; Elvira-Matelot et al., 2016). Here we describe two additional mutants identified during this screen. Based on allelic tests, these two mutants define the *SGS15* locus. In addition to PTGS-deficiency, the *sgs15-1 and sgs15-2* mutants exhibit a late flowering phenotype (Figure 1A). Whole-genome sequencing revealed a deletion in *sgs15-1*, removing part or all of six protein-coding genes (At1g04020, At1g040340, At1g04040, At1g04050, At1g04060, and At1g04080), while *sgs15-2* exhibits a 725 kb inversion on chromosome 1, which disrupts At1g01950 and At1g04080 (Figure 1B). At1g04080 is the only common gene mutated in both *sgs15-1 and sgs15-2*, suggesting that it is responsible for late flowering and S-PTGS defect. At1g04080 was previously described as PRP39a, one of the two Arabidopsis genes encoding a protein homologous to the yeast splicing factor PRP39 (Wang et al., 2007; Kanno et al., 2017). Homozygous *prp39a-1* plants, which harbor a T-DNA insertion in At1g04080 (SAIL_1249A03), exhibited a late flowering phenotype similar to *sgs15-1 and sgs15-2* (Wang et al., 2007; Kanno et al., 2017) and crosses between *prp39a-1* and *sgs15-1 and sgs15-2* yield late flowering F1 plants, suggesting that mutations in At1g04080 are responsible for the late flowering phenotype of *prp39a-1 sgs15-1* and *sgs15-2* mutants.

**Figure 1:**
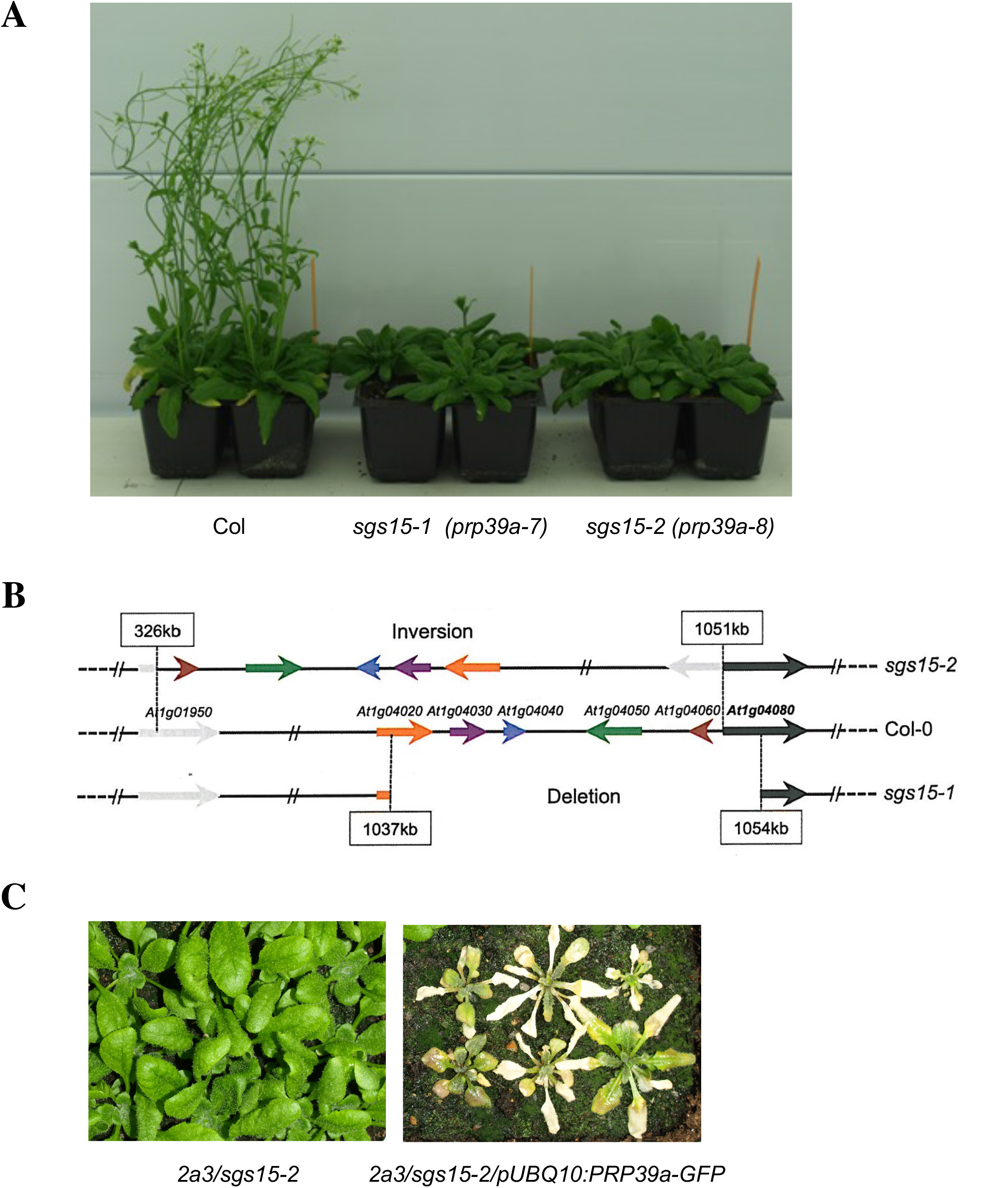
Mutations in PRP39a cause late flowering and suppress S-PTGS. A) Photographs of 40-day-old plants of wild-type Col, *prp39a-7 (sgs15-1)* and *prp39a-8 (sgs15-2)* mutants. B) Structure of the top of chromosome 1 in wild-type Col, *prp39a-7 (sgs15-1)* and *prp39a-8 (sgs15-2)*. A deletion removed the 5’ half of the *PRP39a* gene in *prp39a-7 (sgs15-1)*, while an inversion disrupted the 5’ end of the *PRP39a* gene in *prp39a-8 (sgs15-2)*. C) Photographs of 20-day-old *2a3/prp39a-8 (sgs15-2)* plants, which resist to *NIA* S-PTGS, and *2a3/prp39a-8 (sgs15-2) /pUBQ10:PRP39a* transformants, which undergo *NIA* S-PTGS.

Introduction of the *2a3* locus into the *prp39a-1* mutant by crossing, yield plants that do not undergo *NIA* S-PTGS. Moreover, crosses between *2a3*/*prp39a-1* and *2a3*/*sgs15-1* or *2a3*/*sgs15-2* yield F1 plants that do not undergo *NIA* S-PTGS, indicating that mutations in At1g04080 are also responsible for *NIA* S-PTGS defect. To definitely confirm this, we transformed the *sgs15-2* mutant with a construct carrying genomic sequences of the At1g04080 gene comprised between the ATG and the stop codon under the control of the pUBQ10 promoter and fused to GFP, hereafter referred to as *pUBQ10:PRP39a-GFP*. At first, we transformed *sgs15-2* mutant plants from which the *2a3* locus has been segregated away. Among 30 *sgs15-2/pUBQ10:PRP39a-GFP* transformants, 27 developed like wild-type plants with regards to flowering time, confirming that the deletion of At1g04080 is responsible for the late flowering phenotype of the *sgs15-2* mutant. We also transformed *sgs15-2* mutant plants carrying the *2a3* locus. Among 55 transformants *sgs15-2/pUBQ10:PRP39a-GFP*, 51 exhibited *NIA* S-PTGS (Figure 1C). Altogether, these results indicate that *sgs15-1* and *sgs15-2* are *prp39a* alleles, hereafter referred to as *prp39a-7* and *prp39a-8*, which exhibit a late flowering phenotype, similar to the previously identified *prp39a* mutants (Wang et al., 2007; Kanno et al., 2017). They also indicate that PRP39a participate to PTGS.

### PRP39a localizes in nucleoplasm but is excluded from the nucleolus

We explored where PRP39a is expressed in the cell using transgenic *sgs15-2* mutants carrying a *pUBQ10:PRP39a-GFP* construct that perfectly complements the late flowering and PTGS deficient phenotypes. Confocal analysis revealed a PRP39a-GFP diffuse signal in the nucleoplasm, and excluded from the nucleolus (Figure 2). This pattern resembles that of SmD1, another splicing factor identified through the 2a3 genetic screen (Elvira-Matelot et al., 2016), although SmD1 also associates to nucleocytoplasmic speckles. (Figure 2). Co-localization experiments performed in *N. benthamiana* using XRN2, XRN3 and XRN4, which localize in the nucleolus, nucleoplasm and cytoplasm, respectively, confirmed the nucleoplasmic localization of PRP39a (Figure 2A).

**Figure 2:**
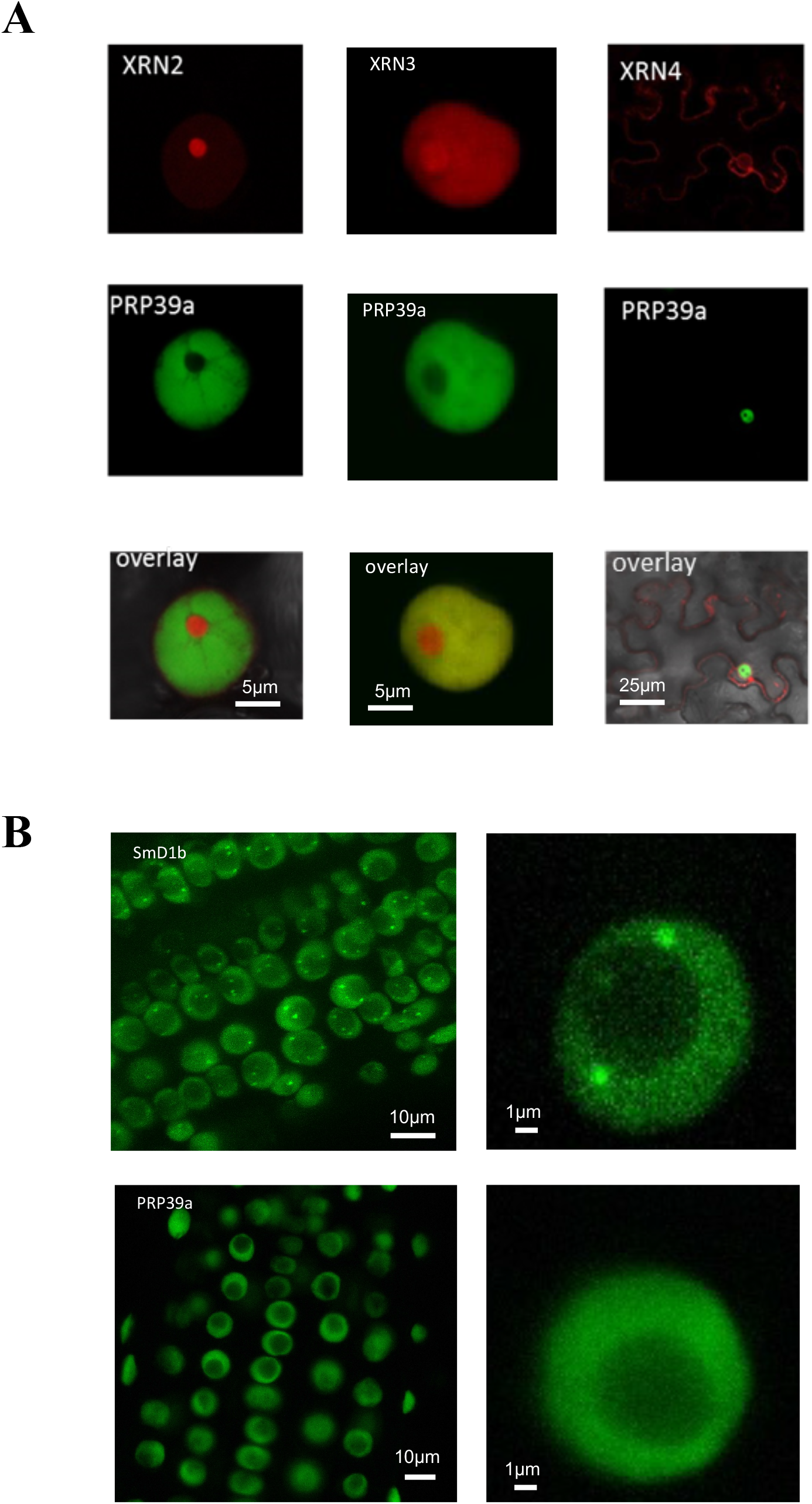
PRP39a subcellular localization. A) Confocal images of *N. benthamiana* leaves infiltrated with *pUBQ10:PRP39a-GFP* reveal the localization of PRP39a in the nucleoplasm. *pUBQ10:XRN2-RFP, pUBQ10:XRN3-RFP* and *pUBQ10:XRN4-RFP* were used as controls for nucleolar, nucleoplasmic and cytoplasmic compartments. B) Confocal images of Arabidopsis root expressing *pUBQ10:PRP39a-GFP or pUBQ10:SmD1b-GFP* reveal distinct nuclear localization

### The abundance and splicing of certain endogenous RNA is differently affected in prp39a and smd1b mutants

In yeast, PRP39 is part of the U1 snRNP complex, which plays important roles in splicing. In Arabidopsis, *prp39a* mutants were previously identified in a forward genetic screen for mutants deficient in pre-mRNA splicing (Kanno et al., 2017). Moreover, Kanno et al., 2017 reported RNA-seq data and analyses of differentially expressed genes (DEGs) and differentially alternatively splicied (DAS) events, confirming the role of Arabidopsis PRP39a in splicing. RNAseq performed on *prp39a* in our lab revealed little overlap with the DEGs identified by Kanno et al 2017, but 220 common DAS (Supplemental Figure S1). Considering that Kanno et al., 2017 used different mutant alleles, developmental stages and DAS analysis methods, this significant overlap between DAS support a major role of PRP39a in AS regulation.

Because *prp39a* and *smd1b* mutants were both recovered from the *2a3* PTGS screen, RNAseq was performed in parallel on *prp39a* and *smd1b* mutants to compare the impact of these two conserved spliceosomal components on RNA splicing and RNA abundance. Both *prp39a* and *smd1b* mutations affect the abundance of numerous transcripts, including mRNAs and a number of non-coding RNAs such as long non-coding RNAs (lncRNA) and natural antisense RNAs (NAT) (Figure 3A). DEGs include intron-less and intron-containing transcripts. The proportion of intron-containing compared to intron-less genes was not significantly different in DEGs than it is in all detected transcripts in the dataset. On the contrary, differentially expressed lncRNAs and differentially expressed NATs were significantly enriched for intron-containing transcript (p<0.001, Fisher exact test) suggesting a potential role of lncRNA splicing in the control of their abundance (Supplemental Figure S2). Although both proteins are splicing regulators, limited overlap was observed between DEGs in both mutants (Figure 3B). Clustering analysis of DEG expression identified cluster of genes with distinct expression patterns in *prp39* and *smd1b* (Figure 3C). Gene ontology enrichment analysis of mRNA clusters (Table 1) showed that genes specifically upregulated in *smd1b* (Cluster 3) are enriched for GO categories associated with defense responses such as response to chitin (p.hyper < 5E-16), response to jasmonic acid (p.hyper < 2.3E-6) and glucosinolate metabolism (p.hyper < 2.5E-5). Gene clusters specifically up-regulated in *prp39a* were enriched for growth and development GO such as auxin response (p.hyper < 2.6E-7), regulation of growth (p.hyper < 4.6E-6) and negative regulation of flower development (p.hyper < 24E-4). Consistently with the late flowering phenotype in the mutant, the master floral regulator *FLC* accumulated in *prp39a* seedlings compared to WT (Supplemental Figure S3A). Opposite to *smd1b*, specifically down-regulated genes (Cluster 5) in *prp39a* were associated with plant defense responses (Response to bacterium, p.hyper < 3.6E-08; immune system process, p.hyper < 3E-5).

**Figure 3:**
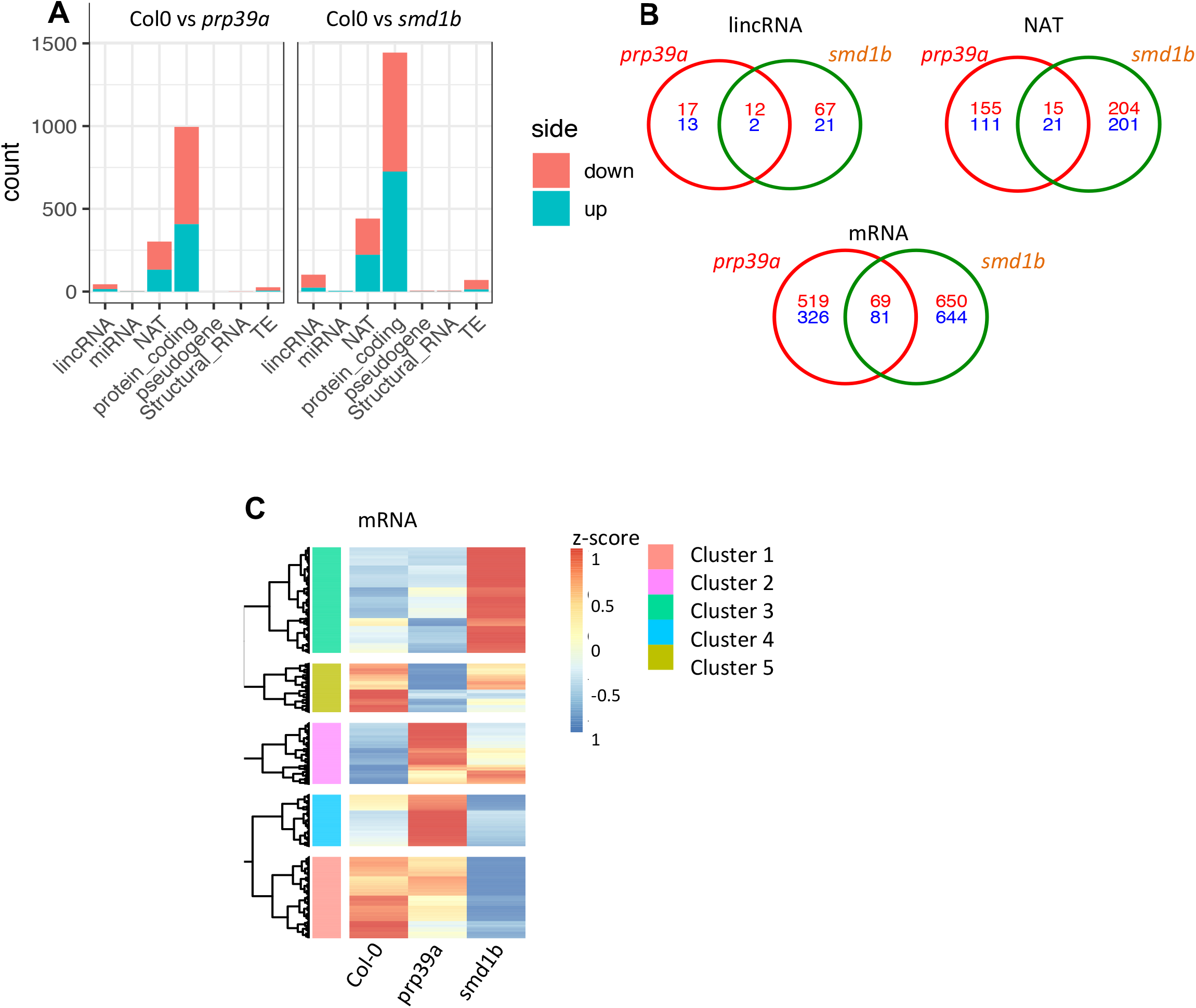
Changes in RNA abundance in *smd1b* and *prp39a* compared to Col-0. A) Number of up and down regulated genes of each class of RNA for *prp39a* and *smd1b* compared to Col-0. B) Overlap of up and down-regulated lincRNA, NAT and mRNA genes in *prp39a* and *smd1b* compared to Col-0 C) Hierarchical clustering of the normalized RNA abundance in Col-0, *prp39a* and *smd1b*. Significant clusters of mRNA genes showing distinct expression profiled are highlighted with colors.

**Table 1.** Gene ontology analysis of gene clusters described in Figure 3C.

| GO TERM | FDR | Cluster ID |
| --- | --- | --- |
| response to chitin | 9.22E-14 | 3 |
| response to wounding | 1.81E-09 | 3 |
| response to hypoxia | 2.53E-09 | 3 |
| ethylene-activated signaling pathway | 0.000216728 | 3 |
| response to jasmonic acid | 0.000391466 | 3 |
| glucosinolate metabolic process | 0.003900072 | 3 |
| phloem or xylem histogenesis | 0.00847591 | 3 |
| auxin response | 3.94E-05 | 4 |
| response to light stimulus | 0.000511594 | 4 |
| regulation of growth | 0.000711629 | 4 |
| water transport | 0.00116582 | 4 |
| negative regulation of flower development | 0.061579308 | 4 |
| response to bacterium | 9.74E-06 | 5 |
| drug transport | 8.08E-05 | 5 |
| response to drug | 0.000491999 | 5 |
| indole-containing compound biosynthetic process | 0.00159862 | 5 |
| iron assimilation by chelation and transport | 0.003719223 | 5 |
| energy derivation by oxidation of organic compounds | 0.004333832 | 5 |
| drug transmembrane transport | 0.005272814 | 5 |
| response to nematode | 0.006465477 | 5 |
| immune system process | 0.008049891 | 5 |

To evaluate the impact of *prp39a* and *smd1b* on mRNA differential splicing, we determined transcript isoforms abundance using the AtRTD2 transcript database (Zhang et al., 2017) and we computed the difference of percentage of spliced-in (ΔPSI) of each possible splicing event genome-wide using SUPPA2 (Trincado et al., 2018). Data showed that *prp39a* had a stronger impact on alternative splicing than *smd1b* when compared to Col-0, with 356 and 142 high confidence DAS (differentially alternatively spliced) genes respectively (ΔPSI >0.2, p.adj < 0.001). Both mutations affected all type of alternative splicing events with intron retention being the most abundant (Figure 4A). Manual evaluation of mRNA-seq read coverage confirmed intron retention events in the *prp39a* background (three examples of AS events are shown in Supplemental Figure S3B). Analysis of DAS genes show that *prp39a* seems to affect preferentially transcript with smaller exons and higher number of introns and that DAS in *prp39a* and *smd1b* have significantly smaller introns that non-DAS transcript (Supplemental Figure S4A). GC content analysis of differential introns and their flanking exons indicated that both mutants preferentially target introns having low GC content in their 3’ flanking exons (Supplemental Figure S4B). As previously presented for transcript abundance (DEGs), both mutations affect splicing of different group of transcripts and DAS genes were largely different from DEGs genome-wide (Figure 4B). To further compare the impact of both mutations on splicing, we compared ΔPSI for each individual intron retention event in both mutants. The analysis shows that Intron Retention (IR) changes were not globally correlated in both mutants (as compared to wild type) except for a small group of introns. Overall, PRP39a and SmD1 appear as AS regulators acting on different splicing targets.

**Figure 4:**
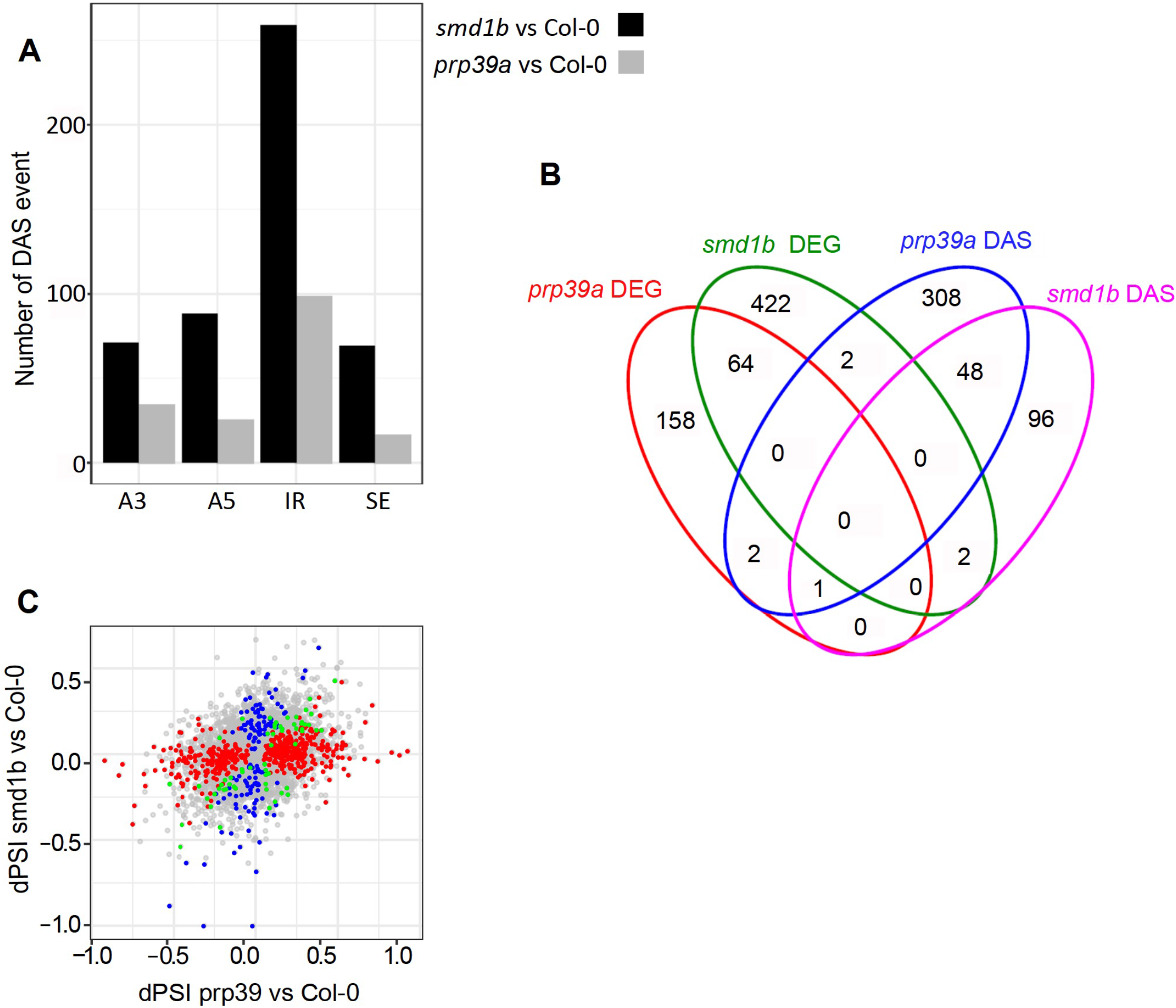
Changes in RNA abundance in *smd1b* and *prp39a* compared to Col-0. A) Number of each class of AS showing significant change in *prp39a* or *smd1b* compared to Col-0. (A3, Alternative 3’ Splice Site; A5 Alternative 5’ Splice Site; IR Intron Retention; SE, Skipped Exon) B) Comparison of DEG and DAS genes in *prp39a* or *smd1b* compared to Col-0. C) Comparison Δpsi changes between *prp39a* vs Col-0 and *smd1b* vs Col-0 for all IR events. Significant changes in *prp39a* vs Col-0, *smd1b* vs Col-0 or both are highlighted in red, blue and green, respectively.

To determine whether changes in splicing or expression of genes involved in RNA silencing could explain the effect of *prp39a* or *smd1b* on PTGS, we compared DEG and DAS with a list of manually curated genes from the literature, which have or could have function in TGS, PTGS or RQC. Only very few DAS or DEG overlapped with this gene set (Supplemental Figure S5, Supplemental Dataset S2) and no key player of TGS, PTGS or RQC was found to be mis-expressed or mis-spliced in *prp39a* or *smd1b*. This further suggests that these proteins affect PTGS in other ways than regulating the splicing or expression of known RNA silencing genes.

### The effect of prp39a on transgene S-PTGS is not specific to intron-containing transgenes but rather depends on the strength of the silencing locus

Reminiscent of the *smd1b* mutation (Elvira-Matelot et al., 2016), the *prp39a* mutations do not totally suppress *NIA* S-PTGS. Indeed, ∼60% of the *2a3/smd1b* plants and ∼40% of either *2a3/prp39a-1, 2a3/prp39a-6* or *2a3/prp39a-7* plants still undergo *NIA* S-PTGS at each generation (Figure 5A). To ensure that S-PTGS is actually compromised in *2a3/prp39a* plants, RNAs were extracted from wild-type Col, *2a3* silenced plants, and *2a3/prp39a, 2a3/smd1b* and *2a3/sgs3* non-silenced plants and hybridized with a *NIA2* probe. Whereas *2a3* plants accumulated *NIA2* siRNAs, *2a3/prp39a* mutants lacked *NIA2* siRNAs and accumulated *NIA2* mRNA, similar to *2a3/smd1b* and *2a3/sgs3* mutants (Figure 5B), indicating that the *prp39a* mutation actually affects *NIA* S-PTGS.

**Figure 5:**
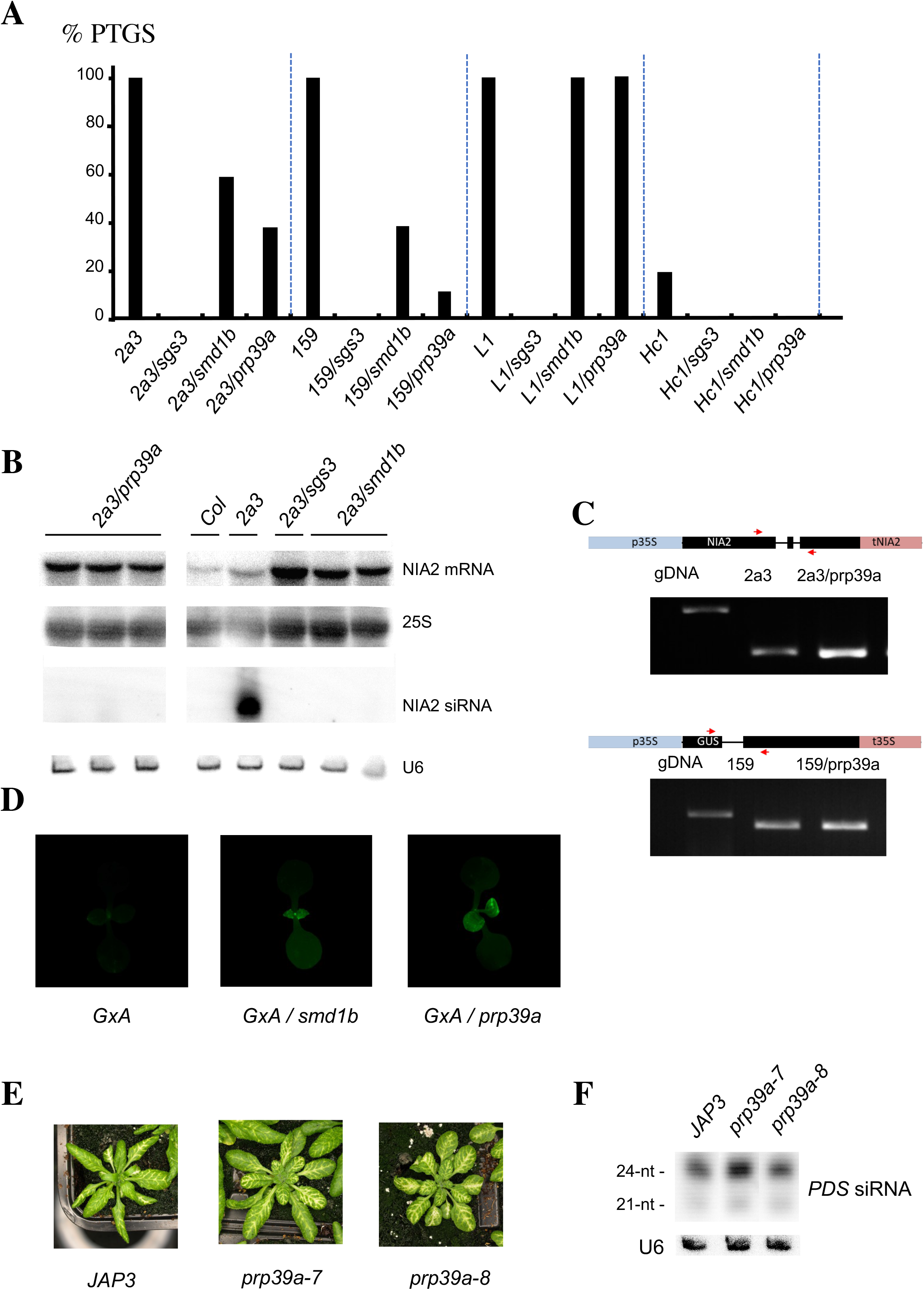
Comparative effect of *prp39a* and *smd1b* mutations on different PTGS systems. A) Percentages of silenced *2a3, L1, Hc1* and *159* plants in the indicated genotypes determined by quantitative GUS activity measurements (n = 96 plants for each genotype). B) RNA gel blot analyses of *NIA* mRNA and siRNAs in the indicated genotypes. *25S* rRNA ethidium bromide staining served as loading controls for high molecular weight RNA blots. *U6* snRNA hybridization served as loading controls for low molecular weight RNA blots. Part of these blots was previously shown in Elvira-Matelot et al, 2016. Original scans can be seen in Supplemental Figure S11. C) Analysis of *NIA2* RNA splicing by RT-PCR using primers spanning an intron. Transgene cassette organization is shown above the gel picture, red arrows mark the relative position of RT-PCR primers. D) Photographs of plants carrying the GxA transgene loci in the indicated genotypes. E) Photographs of plants carrying the JAP3 transgene locus in the indicated genotypes. F) RNA gel blot analyses of *PDS* siRNAs in the indicated genotypes. *U6* snRNA hybridization served as loading control

Similar to the *smd1b* mutant, the *prp39a* mutant was recovered from the S-PTGS genetic screen based on the *2a3* line, but not from the S-PTGS genetic screen based on the *L1* line, which carries a silenced *p35S:GUS* transgene. To test if *prp39a* has an effect on *GUS* S-PTGS, the *L1* locus was introduced into *prp39a-1* by crossing. Similar to *L1/smd1b, L1/prp39a* plants still exhibited *GUS* S-PTGS (Figure 5A), explaining why *smd1b* and *prp39a* mutations were not recovered from the *L1* genetic screen.

The *L1* and *2a3* loci differ by their genomic location and structure, but also by the presence of introns in the *2a3* line. Indeed, the *35S:NIA2* transgene carried at the *2a3* locus derives from a genomic fragment containing the three introns of the Arabidopsis *NIA2* gene. To determine if the basis of the different behavior of the *prp39a* mutation towards *L1* and *2a3* is due to the presence of introns, the *159* locus was introduced into the *prp39a* mutant by crossing. The *159* locus carries the same *p35S:GUS* transgene as *L1* except for the presence of a plant intron within the *GUS* sequence (Vancanneyt et al., 1990). Like *L1*, line *159* exhibits high GUS activity at early stages of development, low GUS activity at later stages in 100% of the population (Figure 5A), and accumulates high levels of *GUS* siRNAs when S-PTGS is triggered (Elvira-Matelot et al., 2016). However, the timing of silencing in line *159* is delayed compared with *L1*, indicating that the *159* locus is a weaker silencing inducer than the *L1* locus. Analysis of *159/prp39a* plants revealed that *GUS* S-PTGS is reduced to 8%, indicating that *prp39a* suppresses *GUS* S-PTGS at this locus, similar to *smd1b*, which reduced *GUS* S-PTGS to 35% (Figure 5A).

To determine if splicing defects where responsible for S-PTGS impairment in *prp39a* plants, we first examined if the endogenous *NIA1* or *NIA2* genes exhibited splicing defects in *prp39a*. Like in *smd1b*, no intron retention or intron skipping was observed in *prp39a* (Supplemental Figure S6). Then, we examined if the *prp39a* mutation causes transgene splicing defects in *159/prp39a* and *2a3/prp39a* plants that could affect S-PTGS. However, the *159* and *2a3* transgenes did not show detectable changes in their splicing patterns (Figure 5C), suggesting that the *prp39a* mutation does not compromise transgene splicing. Thus, S-PTGS impairment in *prp39a* does not appear to result from perturbed splicing of *NIA* endogenous genes or *NIA* or *GUS* transgenes, suggesting that PRP39a acts in S-PTGS independent of its role in splicing, but is dispensable for transgene loci like *L1*, which exhibit very strong S-PTGS.

To test further if *prp39a* only affects S-PTGS at loci that exhibit weak S-PTGS, we first used the *Hc1* locus, which carries the very same *p35S:GUS* transgene as *L1*, but triggers *GUS* S-PTGS in only 20% of the population at each generation whereas *L1* triggers *GUS* S-PTGS with 100% efficiency (Elmayan et al., 1998b; Gy et al., 2007; Martinez de Alba et al., 2011; Martinez de Alba et al., 2015). None of the *Hc1/prp39a* plants triggered *GUS* S-PTGS (Figure 5A), indicating that the *prp39a* mutation also affects S-PTGS of intron-less transgenes. To confirm this, we used the two-components GxA system, where “G” represents a *35S:GFP* transgene and “A” a virus-derived *35S:PVX-GFP* amplicon, which causes *GFP* S-PTGS. Similar to *GxA/smd1b* plants, *GxA/prp39a* plants exhibited GFP expression (Figure 5D), indicating that, similar to *smd1b, prp39a* affects *GFP, GUS* and *NIA2* S-PTGS, but only from weak S-PTGS loci.

Finally, we tested whether *prp39a* affects IR-PTGS. For this, we crossed *prp39a* to the line *JAP3*, which produces *PDS* dsRNA in companion cells of the phloem, resulting in IR-PTGS of the endogenous *PDS* mRNA in a layer of 15 cells around the veins. *JAP3/prp39a* exhibited PDS silencing similar to *JAP3* control plants (Figure 5E), indicating that it has no effect on IR-PTGS, similar to *smd1b* (Elvira-Matelot et al., 2016).

Altogether, these results indicate that PRP39a participates in a step that is specific to S-PTGS. The *prp39a* mutation has a significant effect on most S-PTGS loci (*Hc1, 2a3, GxA, 159*), with the exception of the very strong S-PTGS loci *L1*, indicating that PRP39a facilitates S-PTGS but is not a core component of the S-PTGS pathway, similarly to SmD1 (Elvira-Matelot et al., 2016).

### PRP39b does not appear to play a role in S-PTGS

In Arabidopsis, SmD1 is encoded by two genes, *SmD1a* and *SmD1b*, which act redundantly, but *SmD1b* plays a major role, likely because it is more expressed than *SmD1a* (Elvira-Matelot et al., 2016). Yeast PRP39 has two orthologs in Arabidopsis, At1g04080 and At5g46400, which are referred to as PRP39a and PRP39b, respectively (Supplemental Figure S7A). In accordance with Kanno et al., 2017, we observed that, At5g46400/PRP39b is less expressed than At1g04080/PRP39a (Supplemental Figure S7B). To determine if these two proteins play similar roles in S-PTGS, a T-DNA insertion line in the PRP39b gene (SAIL_157_B08), hereafter referred to as *prp39b-2*, was crossed to lines *159* and *2a3*. Unlike *159/prp39a, 159/prp39b* plants undergo *GUS* S-PTGS with 100%, similar to *159* control plants (Supplemental Figure S7C). Moreover, all *2a3/prp39b-2* plants die from *NIA* S-PTGS as *2a3* control plants. Together, these results suggest that PRP39a, but not PRP39b, plays a role in S-PTGS. This is unlikely due to the simple fact that PRP39a is more expressed than PRP39b but certainly reflects a lack of redundancy between PRP39a and PRP39b, as previously shown for PRP39a splicing activity (Kanno et al., 2017).

### PRP39a does not participate to the endogenous small RNA repertoire

Because many of the PTGS-deficient mutants previously identified in our screen are impaired in components of the cellular machinery producing endogenous siRNAs (Elmayan et al., 1998b; Fagard et al., 2000; Mourrain et al., 2000; Morel et al., 2002; Boutet et al., 2003; Jauvion et al., 2010; Le Masson et al., 2012), we examined the accumulation of representative endogenous small RNAs (miRNAs, ta-siRNAs and p4-siRNAs) in *prp39a* mutants. The accumulation of miR775, TAS1-, TAS2- and TAS3-tasiRNAs, and siR1003 was not affected in *prp39a* mutants (Supplemental Figure S8A). To further determine the impact of *prp39a* on sRNA biogenesis, we performed sRNA sequencing on *prp39a* and Col-0 seedlings. Accumulation of 21- to 24-nt siRNAs from introns, exons, promoter and intergenic regions did not show any global changes in *prp39a* as compared to Col-0 (Supplemental Figure S8B). We then used ShortStack to detect siRNA clusters *de novo* and analyzed their differential expression. The expression pattern of siRNA clusters was strongly correlated between *prp39a* and Col-0 (Supplemental Figure S8C). In addition, among the 891 21-22nt long and the 23179 24nt-long siRNA clusters detected in our dataset, only 19 and 96 were differentially regulated, respectively (Supplemental Figure S8C). Together these data indicate that PRP39a does not generally contribute to the production or distribution of small RNAs genome-wide, but likely affects transgene S-PTGS at a different step.

### PRP39 and SmD1 interplay with distinct nuclear RQC factors

The results presented above suggest that, like SmD1, PRP39a is not essential for IR-PTGS and is not a core component of S-PTGS, but somehow facilitates S-PTGS. Because RQC does not interplay with IR-PTGS (Moreno et al, 2013) but limits S-PTGS by degrading part of the transgene aberrant RNAs that provoke the activation of S-PTGS (Gy et al., 2007; Moreno et al., 2013; Lange et al., 2014; Martinez de Alba et al., 2015), we propose that, like SmD1, the nuclear factor PRP39a facilitates S-PTGS by interplaying with the degradation of transgene aberrant RNAs by the nuclear RQC machinery, thus favoring their transformation in dsRNAs that trigger S-PTGS. To test this hypothesis, *prp39a rqc* double mutants were generated by crossing *159/prp39a* plants with *hen2* and *xrn3* mutants impaired in 3’-to-5’ and 5’-to-3’ degradation of abRNA in the nucleus, respectively. In parallel, *159/smd1b* plants were crossed with *hen2* mutants to complete the analysis previously initiated by crossing *159/smd1b* with *xrn3* (Elvira-Matelot et al., 2016). *GUS* PTGS was restored in *159/prp39a hen2* but not *159/prp39a xrn3* (Figure 6A). In contrast, *GUS* S-PTGS was restored in *159/smd1b xrn3*, but not *159/smd1b hen2* (Figure 6A), suggesting that PRP39 and SmD1 promote S-PTGS by limiting the action of distinct nuclear RQC factors against the aberrant RNAs that are converted to dsRNA by RDR6 to initiate S-PTGS.

**Figure 6:**
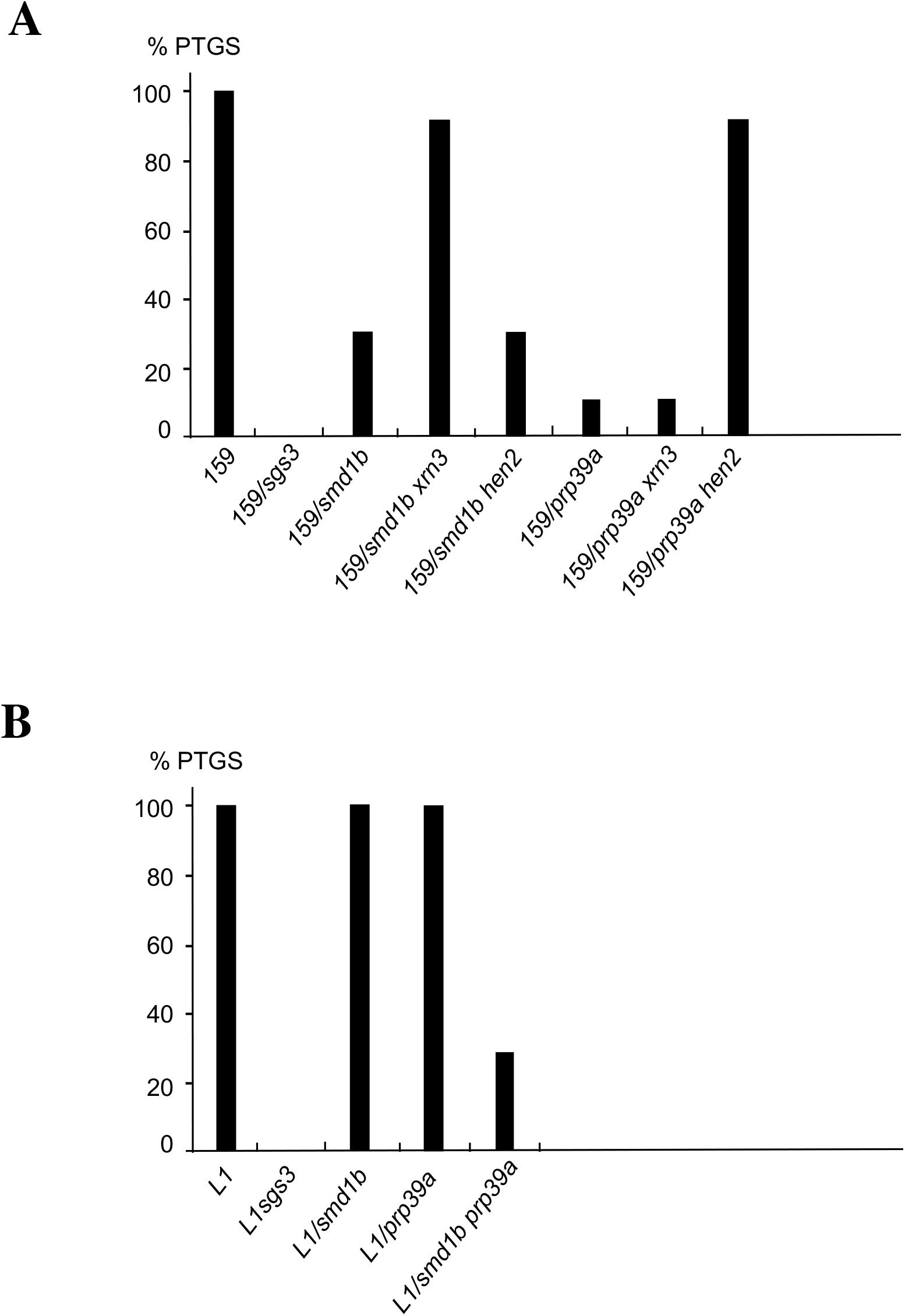
Interplay between *prp39a* and *smd1b* mutations and RQC mutations. A) Distinct RQC mutations suppress the effect of *prp39a* and *smd1b* mutations on S-PTGS. Percentages of plants silenced by S-PTGS in the indicated genotypes were determined by quantitative GUS activity measurements (n = 96 plants for each genotype). B) *prp39a* and *smd1b* have synergistic effect on strong S-PTGS triggered by the L1 locus. Percentages of plants silenced by S-PTGS in the indicated genotypes were determined by quantitative GUS activity measurements (n = 96 plants for each genotype).

### PRP39 and SmD1 have additive effect in promoting S-PTGS

The results presented above indicate that PRP39 and SmD1 promote S-PTGS by distinct means and, as shown for alternative splicing regulation, they define two independent pathways. To test if PRP39 and SmD1 have additive effects, the *L1/prp39a* line, which still undergoes S-PTGS with 100% efficiency was crossed with the *L1/smd1b* line, which also undergoes S-PTGS with 100% efficiency. Analysis of the *L1/prp39a smd1b* double mutant revealed a strong reduction of *GUS* S-PTGS (Figure 6B), supporting the hypothesis of a synergistic effect of these two mutations in promoting S-PTGS.

### PRP39a does not bind to transgene RNA

The absence of a detectable effect of *prp39a* on the splicing of transgene *GUS* and *NIA* RNAs, and the ability of *prp39a* to suppress S-PTGS triggered by intron-less transgenes, suggest that PRP39a facilitates S-PTGS independently of its role in splicing. To determine if, like SmD1, PRP39a directly interacts with transgene RNA, the *159/prp39a* line was transformed with the *pUBQ10:PRP39a-GFP* construct, and complemented transformants that developed like wild-type plants and lacked GUS activity were selected to perform RNA immunoprecipation assays (RIP). The nuclei extract (Input) of the *159* line and one *159/prp39a/pUBQ10:PRP39a-GFP* transgenic line that triggered *GUS* S-PTGS as efficiently as the *159* line were used for RIP using anti-GFP antibodies to detect transgene RNAs bound to PRP39a-GFP. To determine the efficiency of the PRP39-GFP immunoprecipitation in the RIP buffer, we performed a western blot of the input (nuclei extract), unbound and IP (ImmunoPrecipitated) fraction in two *159/prp39a/pUBQ10:PRP39a-GFP* transgenic lines using GFP antibody and an IGG negative control. Results showed no signal in the unbound fraction and a stronger signal in the IP fraction when GFP antibodies were used. On the other hand, IGG IP did not recover any protein signal. Together this showed that PRP39a-GFP was quantitatively recovered during IP and the IP did not produce any detectable background. A *159/smd1b/pUBQ10:SmD1b-GFP* line was included as positive control because SmD1 was previously shown to bind *GUS* RNA (Elvira-Matelot et al., 2016). Specific pairs of primers that amplify *GUS* mRNA (Figure 7) were used to perform Reverse Transcription followed by quantitative real-time PCR on the RIP and the Input samples. As positive controls, we also included three primer sets targeting pre-mRNA transcripts identified as differentially spliced in the *prp39a* RNA-seq analysis. Results showed an enrichment for the three pre-mRNAs targeted by PRP39a, but did not show any enrichment of *GUS* RNA in *159/prp39a/pUBQ10:PRP39a-GFP* plants (Figure 7), indicating that PRP39a does not seem to bind to RNAs produced by silenced transgenes.

**Figure 7:**
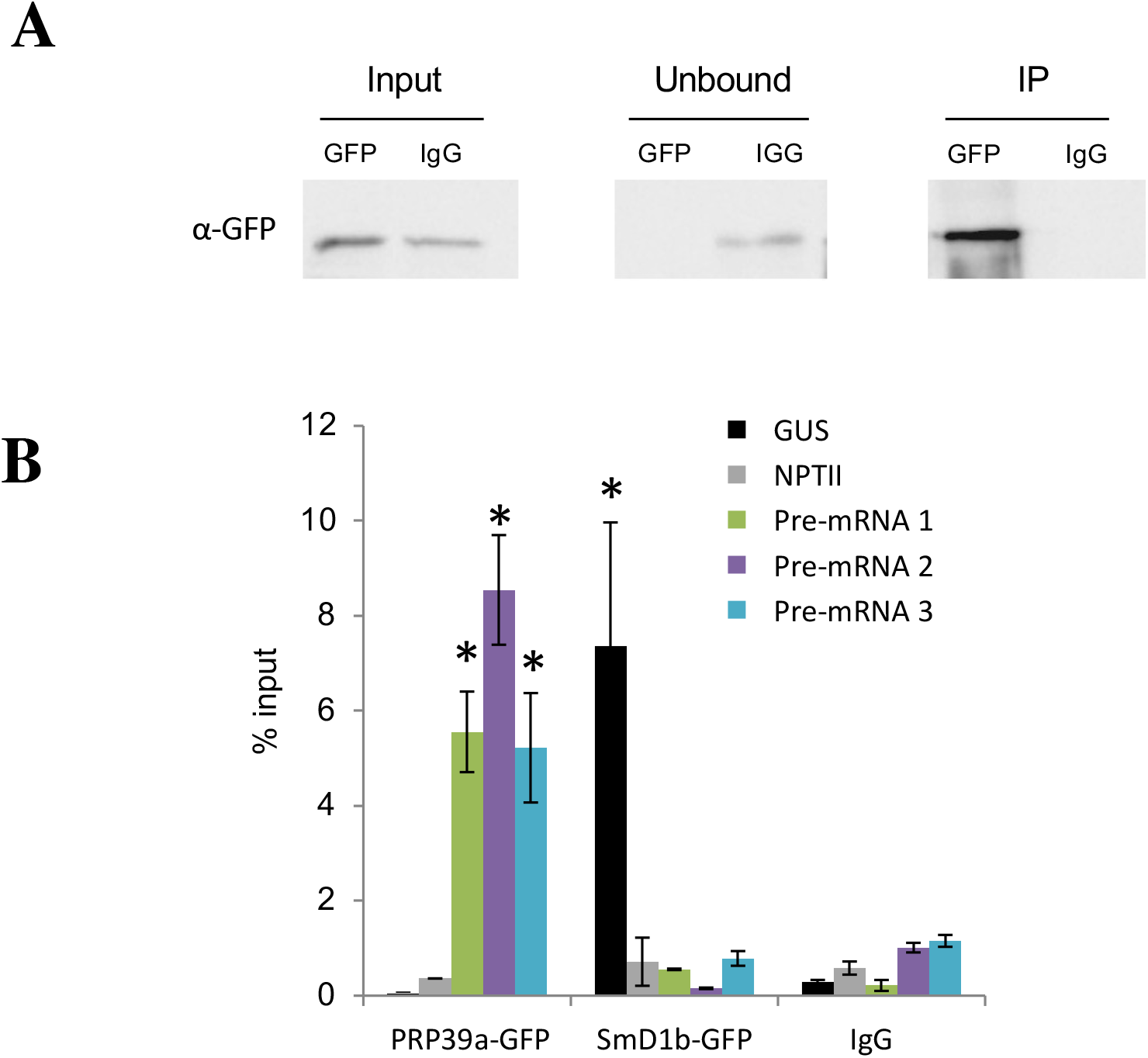
Transgene RNA immunoprecipitation in *159/prp39a/pUBQ10:PRP39a-GFP* and *159/smd1b/pUBQ10:SmD1b-GFP* plants. Lines exhibiting a full restoration of *GUS* S-PTGS were used for immunoprecipitation using GFP antibodies. A) Western blot analysis of the PRP39a-GFP immunoprecipitation, the membranes was probed with GFP antibodies (α-GFP). Input, unbound and IP fraction from immunoprecipitation with GFP antibodies or normal Rabbit IgG was loaded on gel. B) RIP-qRT-PCR analysis using *GUS* and *NPTII* primers. As positive control for PRP39a-GFP RIP, we included three primer sets targeting pre-mRNA transcripts identified as differentially spliced in the *prp39a* RNA-seq analysis. Data represent the mean percentage of input of 3 biological replicates. Error bars show standard error of the mean. Significant enrichment over the IgG RIP was determined using a student-t test (* p.val <0.05)

### PRP39 largely reduce RNA degradation by the nuclear exosome

The results presented above indicate that whereas SmD1 binds to transgene aberrant RNAs to prevent their degradation by the exoribonuclease XRN3 involved in the 5’-to-3’ RNA degradation, PRP39a does not bind to transgene abRNAs but somehow prevents 3’-to-5’ transgene abRNA degradation by the HEN2-containing nuclear exosome. The exact nature of the abRNA that initiate *GUS* S-PTGS is still not known; however, we previously identified an uncapped RNA antisense to *GUS* (referred to as *SUG*). *SUG* RNA is detected in *rdr6*, indicating that it is not an RDR6 product. *SUG* accumulates at higher level in *rdr6* mutants than in wildtype plants, indicating that, like *GUS* mRNA, it is targeted by *GUS* siRNAs. *SUG* accumulates at even higher levels in *rdr6 xrn3 xrn4* mutants, indicating that it is also a target of RQC (Parent et al, 2015). We therefore examined *SUG* accumulation using strand specific qRT-PCR in *159, 159 prp39a, 159 smd1b* and *159 rdr6*. Similar to *159 rdr6, 159 smd1b* accumulated high levels of *SUG* (Supplemental Figure S9). Given the sensitivity of the uncapped *SUG* RNA to XRN-mediated degradation (Parent et al, 2015), this result supports the proposed role of SmD1 in binding transgene abRNA to protect them against 5’-to-3’ degradation (Figure 6A). In contrast, *SUG* level was only slightly increased in *159 prp39a*. This result also is consistent with the fact that PRP39a does not bind transgene RNA (Figure 7), and likely limits the action of the nuclear exosome on aberrant RNAs indirectly.

To test this hypothesis, we examined the accumulation of known HEN2 targets (Lange et al., 2014) in *prp39a*, assuming that, in the absence of PRP39a, these targets should be more efficiently degraded than in wild type plants. Approximately half of the known HEN2 targets (41 out of 97) that we could detect in both RNA-seq dataset were down-regulated in *prp39a* (Figure 8A), supporting this hypothesis. We further examined the expression of a subset of HEN2 targets in *prp39, hen2* and *prp39 hen2* seedlings. As previously shown (Lange et al., 2014), we observed a strong accumulation of short transcripts derived from mRNAs, 3’extension of mRNAs, and incompletely spliced transcripts in *hen2* (Figure 8). In *prp39a*, these transcripts were only marginally affected, but the introduction of the *prp39a* mutation in the *hen2* mutants background strongly affected the accumulation of these exosome targets, indicating that PRP39a has a protecting effect on RNAs targeted by the nuclear exosome.

**Figure 8:**
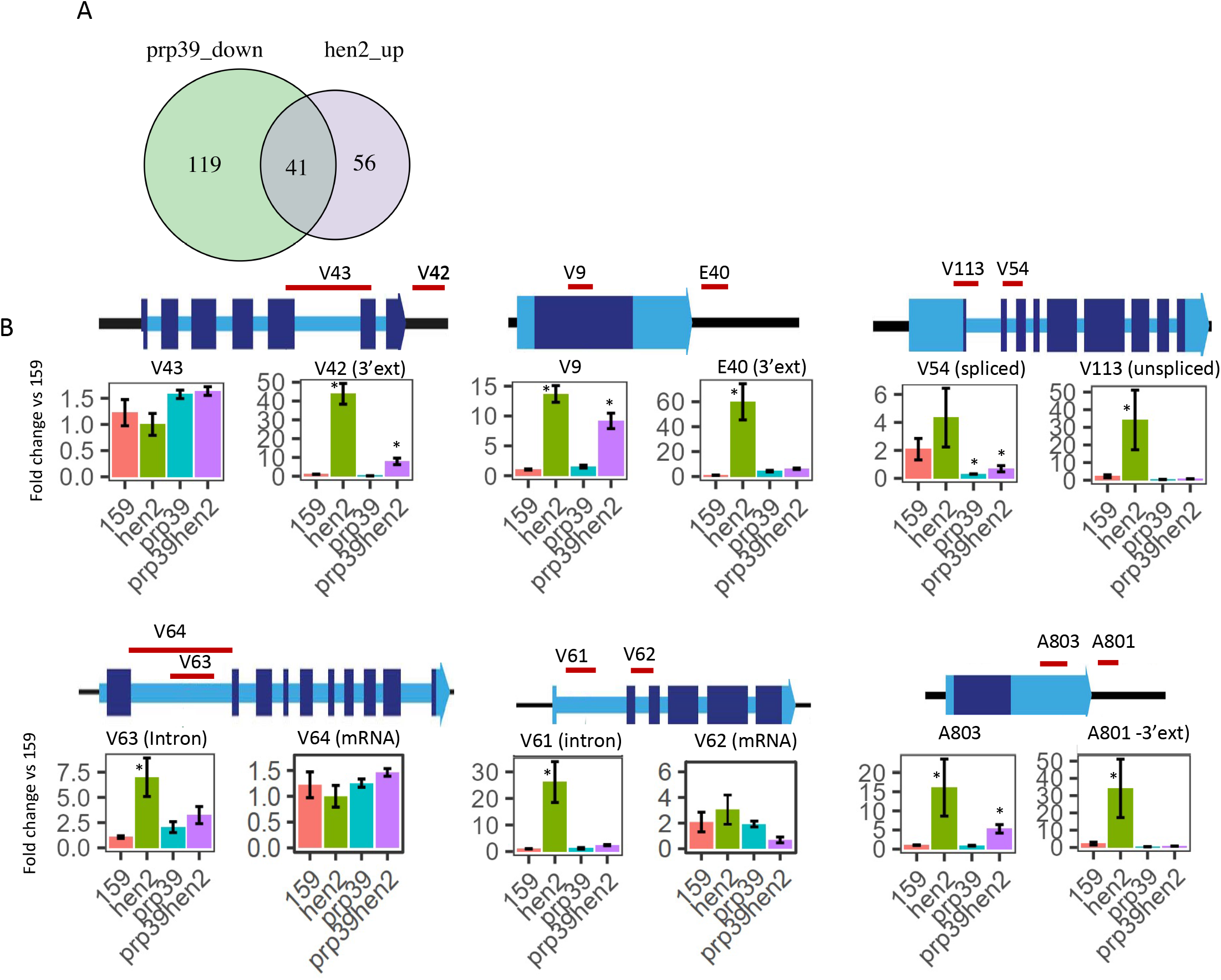
PRP39a generally limits the degradation of HEN2 targets by the nuclear exosome. A) Overlap between down-regulated and up-regulated genes related to WT in *prp39a* and*hen2* mutants, respectively. Only genes detected in both RNA-seq were kept for the analysis. B) Relative expression of selected HEN2 targets determined by qRT-qPCR in WT plants and *prp39a, hen2* and *prp39a hen2* mutants. The red lines show the relative position of qPCR probes on the transcript. Results are mean of 3 biological replicates. Error bars show standard error of the replicates. Significant difference compared to the 159 line was determined using a student-t test (* p.val <0.05) For each gene two pair of primers have been used. One measuring the steady state abundance of the transcript and one measuring the aberrant RNA (3’ extension, unspliced transcript, see Lange et al 2014 for details).

Because the binding of SmD1 to transgene abRNAs likely protects them against 5’-to-3’ degradation (thus explaining why *GUS* S-PTGS is suppressed in *159/smd1b* but restored in *159/smd1b xrn3)*, we examined if impairing SmD1 could have a global effect on endogenous XRN3 targets. For this, we examined the accumulation of a subset of known XRN3 targets (Kurihara et al., 2012) in *smd1b, xrn3* and *smd1b xrn3*. In our conditions, not all known XRN3 targets RNA accumulated in *xrn3*; however, we observed that introducing *smd1b* mutation in the *xrn3* background did not affect the accumulation of the XRN3 targets (Supplemental Figure S10), suggesting that SmD1 does not have a general protective effect on XRN3 substrates, further supporting the hypothesis that PRP39a and SmD1 have distinct actions on PTGS.

## Discussion

RNA interference (RNAi) is a conserved silencing mechanism initiated by dsRNA. In plants, RNAi induced by dsRNA directly resulting from the folding of a self-complementary RNA is referred to as IR-PTGS. However, plant RNAi can also be initiated by (trans)genes that do not directly produce dsRNA, and is referred to as S-PTGS. It is generally assumed that (trans)genes that undergo S-PTGS produce aberrant RNAs (abRNAs) that can be transformed into dsRNA by the RNA-dependent RNA polymerase RDR6. The exact nature of these abRNAs and what makes them specific substrates for RDR6 as well as how they escape degradation by nuclear and cytoplasmic RQC before they reach cytoplasmic siRNA-bodies remains unknown. Recently, we reported that the conserved spliceosome component SmD1b binds transgene RNA and promotes S-PTGS, even when it is triggered by intron-less transgenes (Elvira-Matelot et al., 2016). Because IR-PTGS is not affected in *smd1b* mutants and because S-PTGS is restored in double mutants impaired in SmD1b and certain RQC factors, we proposed that SmD1b protects transgene aberrant RNAs from degradation by certain nuclear RQC pathways. Hence, beside its role in splicing, a spliceosome component can play a role in the interplay between RQC and S-PTGS in the nucleus (Elvira-Matelot et al., 2016).

Here, we describe that another spliceosome component, PRP39a, identified in the same genetic screen for S-PTGS-deficient mutants as SmD1. Like SmD1, PRP39a has dual role in splicing and S-PTGS, and suppresses S-PTGS even when it is triggered by intron-less transgenes (Figure 5). However, SmD1 and PRP39a have distinct roles in both splicing and S-PTGS. RNA-seq analysis of *prp39a* and *smdb1* mutants identified different sets of deregulated RNAs, both at expression level and alternative splicing. Since the relative abundance of alternatively spliced isoforms is mainly influenced by the binding of trans-acting factors such as RNA binding protein and splicing factors on cis-regulatory element in pre-mRNA, it is not surprising that mutants in distinct splicing factors affect splicing of different set of mRNAs. In addition, PRP39 have been shown to be linked to the U1-snRNP component of the spliceosome, which plays critical role in 5’ splice site recognition (Li et al., 2017), whereas SmD1 is a core component of the splicing machinery, suggesting that these two proteins play central but distinct roles in splicing.

Given that *prp39a* and *smdb1* mutants i) do not affect IR-PTGS, ii) affect S-PTGS triggered by intron-less transgenes (Figure 5), and iii) do not modify in the expression or splicing of RQC or PTGS components (Supplemental Figures S3 and S4), PRP39a and SmD1b likely promote a step of PTGS that is specific to the S-PTGS pathway, independently of their role in splicing. Because S-PTGS involves abRNAs that must escape degradation by RQC before being transformed into dsRNA by RDR6, and because PRP39a and SmD1b reside in the nucleoplasm (Figure 2), we examined the interplay between the splicing factors PRP39a and SmD1b and nucleoplasmic RQC factors. Double mutant analyses revealed genetic interactions of PRP39a and SmD1b with distinct nuclear RQC machineries. Indeed, S-PTGS, was restored in *smd1b xrn3* and *prp39a hen2* mutants, but not in *smd1b hen2 and prp39a xrn3* (Figure 6A), suggesting that SmD1b counteracts the action of 5’-to-3’ nuclear RNA degradation, whereas PRP39a counteracts the action of 3’-to-5’ nuclear RNA degradation. RIP-qPCR experiments also revealed that SmD1b binds transgene RNAs whereas PRP39a does not (Figure 7). Therefore, it is likely that the binding of SmD1b to transgene RNA limits its degradation by XRN3, but not by the exosome, whereas PRP39a seems to indirectly limit the degradation of transgene RNAs by the nuclear exosome but not by XRN3. To test whether PRP39a generally counteracts the nuclear exosome, we analyzed the accumulation of known endogenous HEN2 targets in a *prp39a* mutant, and found that PRP39a limits the degradation of almost half of known HEN2 targets (Figure 8), supporting this hypothesis.

How PRP39a limits the activity of the exosome remains to be determined. The structure of the NEXT complex in humans revealed a connection with splicing factors (Falk et al., 2016). Moreover, recent results suggest that the splicing factor SRSF3 may help in the recognition and degradation of certain mRNAs by the nuclear exosome (Mure et al., 2018). Because the Arabidopsis splicing factor PRP39a limits the degradation of certain mRNAs by the nuclear exosome, our results suggest that a connection between splicing factors and RNA degradation also exist in plants, but opposite to that observed in humans. PRP39 is part of the U1 snRNP complex, which beside its role in splicing, is also involved in 3’ end formation by preventing premature transcription termination and cleavage/polyadenylation in vertebrates (Kaida et al., 2010). It is not known if U1 snRNP components participate to 3’ end formation in plants, but Kanno et al., 2017, reported that, in Arabidopsis, PRP39a is highly co-expressed with several cleavage and polyadenylation factors. Given that transgene RNAs lacking a polyA tail are good substrates for RDR6 (Baeg et al., 2017) and highly prone to undergo PTGS (Luo and Chen, 2007), it is tempting to speculate that PRP39a promotes S-PTGS by influencing 3’ end formation and indirectly limiting the activity of the nuclear exosome. Still, PRP39a does not fully prevent the action of the nuclear exosome. Whether the limited effect of the *prp39a* mutation on the accumulation of nuclear exosome targets is due to specific targeting of PRP39a to certain genes only or to the redundant activity of PRP39b remains to be determined. However, PRP39b is expressed at a lower level than PRP39a, and the single *prp39b* mutation has no effect on S-PTGS (Supplemental Figure S5). Moreover, PRP39a and PRP39b were previously shown to not act redundantly in Arabidopsis (Kanno et al., 2017). Therefore, PRP39a may be the only plant PRP39 protein interplaying between splicing, RQC and PTGS.

In conclusion, we propose that PRP39a and SmD1b promote S-PTGS by limiting RQC-mediated degradation of transgene aberrant RNAs using two distinct but complementary manners (Figure 9). SmD1b binds to transgene abRNAs, thus limiting their degradation by XRN3, whereas PRP39a does not bind transgene RNAs but indirectly limits the action of the nuclear exosome, and consequently the degradation of transgene abRNAs. Supporting this hypothesis, a *prp39a smd1b* double mutant exhibits enhanced synergistic suppression of S-PTGS, likely because XRN3 and the nuclear exosome degrade transgene aberrant RNAs more efficiently when both PRP39a and SmD1b are absent. Nevertheless, S-PTGS still occurs to some extent in a *prp39a smd1b* double mutant, indicating that additional nuclear factors are at work to prevent the complete degradation of transgene aberrant RNAs by RQC.

**Figure 9:**
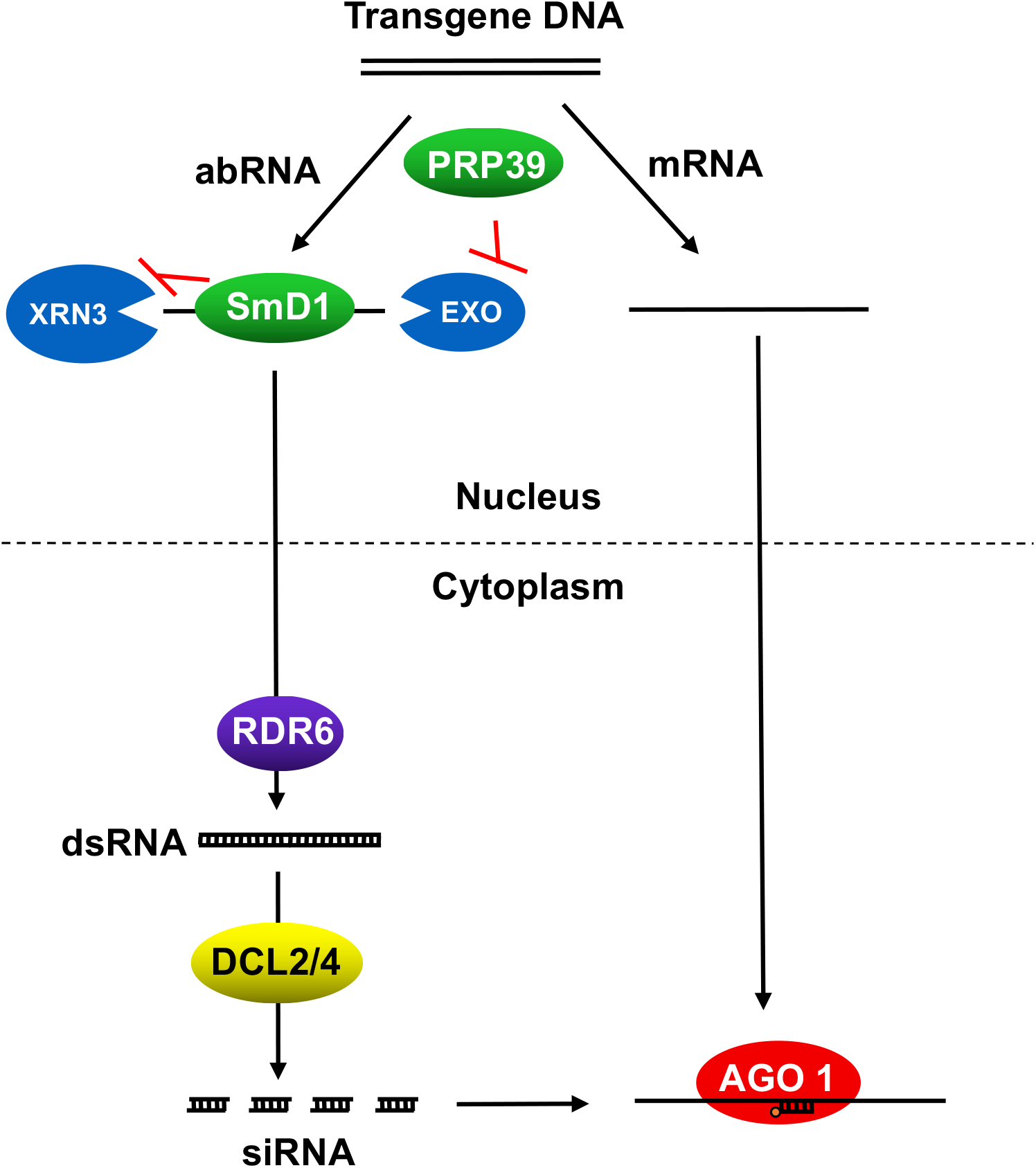
Tentative model for PRP39a and SmD1b action in S-PTGS. Transgene loci that undergo S-PTGS likely produce aberrant RNAs (abRNA). Degradation of these abRNAs by nuclear RQC 5’-to-3, and 3’-to-5’ RNA degradation pathways limits the amount of abRNAs that enter cytoplasmic siRNA-bodies where RDR6 transform them into dsRNA that are processed by DCL2 and DCL4 into siRNAs that are loaded on AGO1 to promote the degradation of regular mRNA. By binding to transgene abRNAs, SmD1b likely limits the binding of XRN3 and limits 5’-to-3 degradation without altering the nuclear exosome activity. In contrast, PRP39a does not bind transgene abRNAs, but somehow limits the action of HEN2 or other components of the nuclear exosome, without altering the activity of XRN3 on these abRNAs. In the absence of PRP39a and SmD1b, transgene abRNAs are more sensitive to 3’-to-5’ and 5’-to-3’ degradation, respectively, thus reducing the probability that a sufficient amount of abRNAs reach siRNA-bodies to activate S-PTGS.

## Methods

### Plant material and growth conditions

All *Arabidopsis thaliana* plants are in the Columbia accession. Transgenic lines *2a3, 159, L1* and *Hc1*, and mutants *hen2 (sop3-2), sgs3-1, smd1b-1, xrn3-3* have been previously described (Elmayan et al., 1998a; Mourrain et al., 2000; Gy et al., 2007; Moreno et al., 2013; Lange et al., 2014; Elvira-Matelot et al., 2016). The T-DNA insertion mutants *prp39a-1* (SAIL_1249A03) and *prp39b-2* (SAIL_157_B08) were obtained from NASC (Alonso et al., 2003). Plants were grown on Bouturage media (Duchefa) in standard long-day conditions (16 hours light, 8 hours dark at 20-22°C) and transferred to soil after two weeks and grown in controlled growth chambers in standard long-day conditions.

### Plasmid constructs

The *pUBQ10:PRP39a-GFP* construct was generated using the Gateway technology (Invitrogen) as follows. The PRP39a genomic fragment was amplified from the first ATG to the penultimate codon to omit the stop codon and cloned into the pENTR/D TOPO vector using pENTR/D TOPO cloning kit (Invitrogen). PRP39a was then transferred into the plant expression vector pB7FWG2 (Karimi et al., 2002) using the LR clonase II kit (Invitrogen). The *p35S:XRN2-RFP, p35S:XRN3-RFP and p35S:XRN4-RFP* constructs have been described previously (Lorkovic et al., 2008; Moreno et al., 2013).

### Arabidopsis transformation and Nicotiana benthamiana agro-infiltration

Agrobacterium strains carrying plasmids of interest were grown overnight at 28°C in 3 ml LB medium containing the appropriate antibiotics to a final OD600 between 1 and 2. For Arabidopsis transformation, the bacteria were pelleted and resuspended in 300 ml of infiltration medium (5% sucrose, 10mM MgCl2, 0,015% silwet L-77) to a final OD600 of 1, which was used for floral dipping. For *Nicotiana benthamiana* agro-infiltration, the bacteria were pelleted and resuspended in 1 ml of infiltration medium (10 mM MgCl_2_, 10 mM MES pH 5.2, 150 mM acetosyringone) to a final OD600 of 0.1. The solution containing the bacteria was injected into the abaxial side leaves using a 1-ml syringe and samples were observed in a confocal microscope 3 days after infiltration.

### Imaging and image analysis

After agro-infiltration, fluorescent cells were imaged by confocal microscopy (Leica TCS SP2, Leica Microsystem, Wetzlar, Germany) with excitation at 488 nm and fluorescence emission signal between 495 and 530 nm for GFP fusions, and excitation at 543 nm and emission signal between 555 and 620 nm for DsRed or RFP fusions. The Leica confocal software was used for image acquisition and for the quantification of fluorescence profiles. Sequential scans were performed when necessary. Spectral profiles were calculated for five cells. Data processing was performed using ImageJ (http://rsbweb.nih.gov/ij/).

### RNA extraction and RNA gel blot analysis

For RNA gel blot analyses, frozen tissue was homogenized in a buffer containing 0.1 M NaCl, 2% SDS, 50 mM Tris-HCl pH 9.0, 10mM EDTA pH 8.0 and 20 mM beta-mercaptoethanol and RNAs were extracted two times with phenol and recovered by ethanol precipitation. To obtain high molecular weigh (HMW) RNA fraction, resuspended RNAs were precipitated overnight in 2M LiCl at 4°C and recovered by centrifugation. For low molecular weigh (LMW) RNA analysis, total RNA was separated on a 15% denaturing PAGE gel, stained with ethidium bromide, and transferred to nylon membrane (HybondNX, Amersham). LMW RNA and U6 hybridizations were at 50°C with hybridization buffer containing 5X SSC, 20mM Na_2_HPO_4_ pH 7.2, 7% SDS, 2X Denhardt’s Solution and denatured sheared salmon sperm DNA (Invitrogen). HMW RNA hybridization was at 37°C in PerfectHyb Plus buffer (Sigma). Blots were hybridized with a radioactively labeled random-primed DNA probes for *GUS* mRNA and *GUS* siRNAs, and an end-labeled oligonucleotide probe for U6 detection.

### RT-PCR analysis

Total RNA was prepared from plantlets at different developmental stages using Trizol (Invitrogen). The DNAse1 treatment (Thermo Scientific) was performed according to the manufacturer’s protocols. For reverse transcription with SuperScriptII (Invitrogen), 2 μg of total DNAse-treated RNA was used. One microliter of the resulting cDNA solution was used for RT-PCR or RT-qPCR analyses. The latter was done using standard protocols and a complete list of RT-qPCR primers is available in Supplemental Table 1. Each cDNA sample was precisely calibrated and verified for two constitutive genes, AT1G13320 and AT4G26410 (Czechowski et al., 2005). For RT-PCR, the amplification was performed as follows: one cycle of 4 min at 98°C, 26 cycles of 30 s at 98°C, 30 s at 59°C, and 1 min at 72°C. The products were separated on agarose gel and stained with ethidium bromide. RT-qPCR was performed using a Roche Light Cycler 480 standard protocol (40 cycles, 60°C annealing).

### RNA Immunoprecipitation

Eleven-day-old plants grown in Petri dishes were irradiated three times with UV using a Crosslinker CL-508 (Uvitec) at 0.400 J/cm^2^. Briefly, fixed material was ground in liquid nitrogen, homogenized, and nuclei isolated and lysed according to (Gendrel et al., 2005).RNA IP was basically performed as described by (Carlotto et al., 2015). The nuclei extract (Input) was used for the immunoprecipitation performed by Direct ChIP Protocol of the Diagenode IP-Star SX-86 Compact robot, using 50 μl of Dynabeads – Protein A (Novex 10008D; Life Technologies, USA) and anti-GFP (632381; Clontech, USA) antibodies. Beads were washed twice for 5 min at 4°C with Wash Buffer 1 (150 mM NaCl, 1% Triton, 0.5% NP-40, 1 mM EDTA, 20 mM Tris-HCl pH7.5) and twice with Wash Buffer 2 (20 mM Tris-HCl pH 8) instead of the Diagenode assigned buffers, and finally resuspended in 100 μl Proteinase K buffer (100mM Tris-HCl pH 7.4, 50 mM NaCl, 10 mM EDTA). After Proteinase K (AM2546; Ambion, USA) treatment, beads were removed with a magneto, and the supernatants were transferred to a 2-ml tube. Each RNA sample was extracted from 800 μl (8 IP-Star tubes) of RNA IP product using 1 ml of TriReagent (Sigma-Aldrich T9424, USA) as indicated by the manufacturer. 80 μl of nuclei extracts were used for Input RNA extraction. The IP and the Input samples were treated with DNase, and random hexamers were used for subsequent RT. Quantitative real-time PCR reactions were performed using specific primers. Results were expressed as a percentage of cDNA detected after IP, taking the Input sample as 100%.

### GUS extraction and activity quantification

GUS protein was extracted and GUS activity was quantified as described before (Gy et al., 2007) from cauline leaves of flowering plants by measuring the quantity of 4-methylumbelliferone product generated from the substrate 4-methylumbelliferyl-b-D-glucuronide (Duchefa) on a fluorometer (Fluoroscan II; Thermo Scientific).

### Transcriptome analysis by RNA sequencing

Total RNA was extracted using RNeasy plant mini kit (Qiagen) from whole 14-day-old Col-0, *prp39a-1* (SAIL_1249A03), *smd1b-1* (Elvirat-Matelot et al., 2016) and Col-0 plants grown on ½ MS medium. Three independent biological replicates were produced per genotype. For each biological repetition and each point. After RNA extraction, polyA RNAs were purified using Dynabeads mRNA direct micro kit (Ambion). Libraries were constructed using the Tru-Seq Stranded mRNA Sample Prep kit (Illumina®). Sequencing was carried out at the POPS Transcriptomic Platform, Institute of Plant Sciences Paris-Saclay in Orsay, France. The Illumina HiSeq2000 technology was used to perform paired-end 100-bp sequencing. A minimum of 30 millions of paired-end reads by sample were generated. RNA-seq preprocessing included trimming library adapters and quality controls with Trimmomatic (Bolger et al., 2014). Paired-end reads with Phred Quality Score Qscore > 20 and read length > 30 bases were kept, and ribosomal RNA sequences were removed with SortMeRNA (Kopylova et al., 2012). Processed reads were aligned using Tophat2 with the following arguments : --max-multihits 1 -i 20 --min-segment-intron 20 --min-coverage-intron 20 --library-type fr-firststrand --microexon-search -I 1000 --max-segment-intron 1000 --max-coverage-intron 1000 --b2-very-sensitive. Reads overlappings exons per genes were counted using the FeatureCounts function of the Rsubreads package using the GTF annotation files from the Araport11 repository (https://www.araport.org/downloads/Araport11_Release_201606/annotation/Araport11_GFF3_genes_transposons.201606.gff.gz). Significance of differential gene expression was estimated using DEseq2 (Love et al., 2014), and the FDR correction of the p-value was used during pairwise comparison between genotypes. A gene was declared differentially expressed if its adjusted p-value (FDR) was ≤ 0.01 and its absolute fold change was ≥ 1.5. Hierarchical clustering analysis was performed in R on scaled normalized read count data. Heatmap was plotted using pheatmap package (https://cran.r-project.org/web/packages/pheatmap/index.html). Gene ontology enrichment analysis was done using the systemPipeR package (Backman and Girke, 2016). Transcript level quantification was performed using pseudo-alignment counts with kallisto (Bray et al., 2016) on AtRTD2 transcripts sequences (https://ics.hutton.ac.uk/atRTD/RTD2/AtRTDv2_QUASI_19April2016.fa) with a K-mer size of 31-nt. Differential AS events in the AtRTD2 database were detected using SUPPA2 with default parameters (Trincado et al., 2018). Intron length, GC content was performed using in-house R script.

### Small RNAseq library construction, sequencing and analysis

Small RNA (sRNA) libraries were constructed from 1µg of total RNA treated with DNaseI (Thermo Scientific) using the NEBNext® Multiplex Small RNA Library Prep Set for Illumina® kit (New England Biolabs) according to manufacturer’s instructions. Libraries were sequenced on a NextSeq 500 Sequencing System (Illumina) using 75-nt single-end reads. The sequencing adapters were removed using cutadapt (v2.10) and sequence matching Arabidopsis ribosomal or transfer RNA were discarded using bowtie (v1.3.1). The reads of length 19 to 25nt, were then mapped on TAIR10 genome with the help of ShortStack (v3.8.5) without mismatch (--mismatches 0), keeping all primary multi-mapping (--bowtie_m all) and correcting for multi-mapped reads according to the uniquely mapped reads (--mmap u). For each gene annotation in Araport11 (coding genes, non-coding RNA and miRNA precursor), 19-nt to 25-nt long read density was counted with ShortStack. sRNA accumulation in each library was normalized using the median of ratio of the corresponding miRNA counts inside DESeq2 (v1.34.0). Differential sRNA accumulation and log2 fold changes between genotypes were computed using DESeq2. FDR correction of the p-value was used.

## Acknowledgements

We thank Hervé Ferry and Philippe Maréchal for plant care, Florian Bardou for his initial contribution to this work, and Taline Elmayan for fruitful discussion. This work was supported by the Agence Nationale de la Recherche ANR-16-CE12-0032 (to HV and MC).

## Authors contributions

HV and MC designed the experiments. All authors contributed to the production and analysis of the results. HV, JB and MC wrote the paper.

## Accession Numbers

Gene sequence data from this article can be found in the GenBank/EMBL libraries under the following accession numbers: PRP39a (At1g04080), PRP39b (At5g46400). RNAseq data generated in this study are accessible through NCBI’s Gene Expression Omnibus (GSE158145).

## Supplemental Data

**Supplemental Figure S1 supporting Figure 3: Overlap between our mRNAseq analysis of *prp39a* and Col-0 and that of Kanno et al**., **2017**

A) overlap between DEG. B) overlap between DAS.

**Supplemental Figure S2 supporting Figure 3: Analysis of transcript classes affected in *prp39a* and *smd1b* mutants compared to Col-0**.

A) Number of intron-containing and intron-differentially expressed genes within each class of transcript (Structural RNA = tRNA and rRNA)

B) Ratio of intron vs intron-less transcripts within each class differentially expressed or detected genes.

**Supplemental Figure S3 supporting Figure 3:** A, Relative mRNA abundance fold change calculated from polyA RNA-seq of FLC (AT5G10140) in smd1b and prp39a as compared to Col-0. (* : FDR<0.05). B, : Normalized mRNA-seq read coverage on three genes showing signifcantly retained introns in prp39a as compared to Col-0. Red boxes show the position of retained introns detected by SUPPA2.

**Supplemental Figure S4 supporting Figure 4: Analysis of exon-intron properties in DAS genes from in *prp39a* and *smd1b* mutants compared to Col-0**.

**A)** Mean introns length, exons length and mean number of introns per DAS genes in *prp39a* and *smd1b* compared to Col-0.

**B)** GC content of flanking 5’ and 3’ exons and introns showing significant intron retentions in *prp39a* and *smd1b* compared to Col-0 (“*prp39a* vs Col-0 introns diff”, “*smd1b* vs Col-0 introns”) compared to non-retained introns (“Introns non diff”)

**A**, Significant differences were calculated with an ANOVA and post hoc Tukey HSD test using

**A**, non-DAS genes (Not DAS) and **B**, non-retained introns (“Introns non diff”) as reference group

**Supplemental Figure S5 supporting Figure 4: Differential expression and splicing of genes encoding silencing and RQC components in *prp39a* and *smd1b* mutants compared to Col-0**.

Overlap of DEG and DAS genes in *smdb1* and *prp39a* compared to Col-0 among genes curated from the literature whose function in TGS, PTGS or RQC has been experimentally demonstrated (listed in Supplemental Dataset S2).

**Supplemental Figure S6 supporting Figure 4: *NIA1* and *NIA2* expression profile in *prp39a* and *smd1b* mutants compared to Col-0**.

RNAseq analysis of *NIA1* and *NIA2* expression in *prp39a* and *smd1b* compared to Col-0 does not reveal defects in intron splicing.

**Supplemental Figure S7: Characterization of the *PRP39b* gene**

A) Alignment of PRP39a and PRP39b proteins. Conserved amino acids are boxed in black and similar amino acids are boxed in grey.

B) ATH1 array expression profiles of *PRP39a* and *PRP39b* genes.

C) Effect of *prp39a* and *prp39b* mutations on S-PTGS triggered by the *159* or *2a3* locus. The percentage of plants showing *GUS* S-PTGS n the indicated genotypes was determined by quantitative GUS activity measurements (n = 96 plants for each genotype). The percentage of silenced plants showing *NIA* S-PTGS in the indicated genotypes was determined by scoring the number of chlorotic plants (n = 96 plants for each genotype).

**Supplemental Figure S8 supporting Figure 5: Effect of *prp39a* mutations on the accumulation of endogenous small RNAs**.

A. RNA gel blot analysis of representative endogenous small RNAs in wild-type Col, *prp39a-7 (sgs15-1)* and *prp39a-8 (sgs15-2)* mutants. *U6* snRNA hybridization served as loading control. B Fraction of 19-nt to 25-nt long siRNA-seq reads mapped to the whole CDS, exons, 3’ and 5’Untranslated Region (3’UTR, 5’UTR), Transcripts, introns and Gene promoters (1kb upstream). C, Normalized abundance of 21-22nt and 24nt siRNA reads on siRNA clusters detected by ShortStack. Red dots mark clusters which show a statistically significant difference of siRNA abundance (DESeq2, FDR <0.01, 3 bioreps).

**Supplemental Figure S9 supporting Figure 5: Accumulation of transgene aberrant RNA SUG measured by strand specific RT-qPCR in plants *159, 159 prp39a, 159 smd1b, 159 sgs3***.

Results are mean of 3 biological replicates. Error bars show standard error of the replicates. Significant difference compared to the 159 line was determined using a student-t test (* p.val <0.05). Primers and the strand specific PCR strategy is described in Parent et al., 2015

**Supplemental Figure S10 supporting Figure 8: SmD1b does not generally limits the degradation of XRN3 targets**.

Relative expression of selected XRN3 targets (as defined in Kurihara et al., 2012) determined by qRT-qPCR in 159 plants and 159 *smd1b,159 xrn3* and 159 *smd1b xrn3*. Results are mean of 3 biological replicates. Error bars show standard error of the replicates

**Supplemental Figure S11: original blots for Figure 5B**

RNA gel blot analyses of *NIA* mRNA and siRNAs in the indicated genotypes. *25S* rRNA ethidium bromide staining served as loading controls for high molecular weight RNA blots. *U6* snRNA hybridization served as loading controls for low molecular weight RNA blots. The part of the blots previously shown in Elvira-Matelot et al, 2016 are indicated in blue, while the parts of the blots shown in Figure 5B are indicated in red.

**Supplemental Dataset S1: Primers used in this study**

**Supplemental Dataset S2: List of curated PTGS and RQC genes analyzed in Supplemental Figure S5**.

DEG and DAS genes between *prp39a-7* and Col-0 were compared to a set of genes manually curated from the literature to be involved in PTGS, or RQC. Column “prp39_vs_col”,”smd1_vs_col” and “PTGS or RQC related genes”. 0 and 1 means presence /absence of the the gene in the considered gene set. When available the gene Name in indicated in the Name column and the biochemical pathway is described in the pathway column. FDR, False Discovery Rate. dPSI (percent spliced index difference). Event show the unique ID of the AS event as defined by SUPPA2.

